# Metabolic endotoxemia is dictated by the type of lipopolysaccharide

**DOI:** 10.1101/2021.07.13.452270

**Authors:** Fernando F. Anhê, Nicole G. Barra, Joseph F. Cavallari, Brandyn D. Henriksbo, Jonathan D. Schertzer

**Affiliations:** Department of Biochemistry and Biomedical Sciences, McMaster University, 1200 Main St. W., Hamilton, Ontario, Canada L8N 3Z5; Farncombe Family Digestive Health Research Institute, McMaster University, 1200 Main St. W., Hamilton, Ontario, Canada L8N 3Z5; Centre for Metabolism, Obesity and Diabetes Research, McMaster University, 1200 Main St. W., Hamilton, Ontario, Canada L8N 3Z5

**Author notes:** **Corresponding Author:** Jonathan D. Schertzer, Biochemistry and Biomedical Studies, McMaster University, 1200 Main Street West, HSC 4H19, Hamilton, ON L8S 4K1.

**Keywords:** Immunometabolism, Inflammation, Obesity, Diabetes, Microbiota, Microbiome

## Abstract

Lipopolysaccharides (LPS) can promote metabolic endotoxemia, which is considered inflammatory and metabolically detrimental based on Toll-like receptor (TLR)4 agonists such as *Escherichia coli*-derived LPS. LPS from certain bacteria antagonize TLR4 yet contribute to endotoxemia measured by Endotoxin Units (EU). We found that *E. coli* LPS impaired gut barrier function and worsened glycemic control in mice, but equal doses of LPS from other bacteria did not. Matching the LPS dose from *R. sphaeroides* and *E. coli* by EU revealed that only *E. coli* LPS promoted dysglycemia, adipose inflammation, delayed intestinal glucose absorption, and augmented insulin and GLP-1 secretion. Metabolically beneficial endotoxemia promoted by *R. sphaeroides* LPS counteracted dysglycemia caused by an equal dose of *E. coli* LPS and promoted insulin sensitivity in obese mice. The concept of metabolic endotoxemia should be expanded beyond LPS load (EU) to include LPS characteristics, where the balance of deleterious and beneficial endotoxemia regulates host metabolism.

**Highlights:**

- Type of LPS dictates gut barrier function, inflammation, insulin, GLP-1, intestinal glucose absorption and blood glucose
- Endotoxin Units (EU) do not reflect how LPS influences blood glucose or hormones
- LPS derived from certain types of bacteria are insulin sensitizers
- *R. sphaeroides* LPS promotes metabolically beneficial endotoxemia
- LPS characteristics dictate metabolically beneficial versus deleterious endotoxemia

## Introduction

Chronic low-grade compartmentalized inflammation is a key aspect of obesity-related insulin resistance and dysglycemia (Hotamisligil, 2006; McPhee and Schertzer, 2015). Blood glucose levels and gut bacteria interact to dictate host metabolic outcomes and bacterial pathogenicity (Anhê et al., 2020a). Lipopolysaccharides (LPS) are a prolific example of an inflammatory bacterial trigger that is modified by obesity and diet. Elevated circulating LPS upon obesogenic feeding (i.e. metabolic endotoxemia) can contribute to metabolic inflammation and dysglycemia (Cani et al., 2007, 2008). Metabolic endotoxemia has been commonly modelled using LPS derived from *Escherichia coli*, which promotes inflammation and metabolic defects via Toll-like receptor (TLR)4 agonism (Amar et al., 2011; Lyte et al., 2016).

The current concept of metabolic endotoxemia does not account for bacterial strain-specific variation in LPS structure and consequent changes in inflammation and metabolism via Myeloid differentiation protein (MD)-2 and TLR4 (Anwar et al., 2015). The polysaccharide region of LPS is anchored in the outer bacterial membrane by lipid A (or endotoxin). Hexa-acylated lipid A generally possesses high TLR4 activation capacity, whereas under-acylated LPS can dose-dependently antagonize TLR4 (Coats et al., 2005; Erridge et al., 2002). Many studies on metabolic endotoxemia have focused on potent hexa-acylated LPS derived from *E. coli*. It is also common to report metabolic endotoxemia in terms of endotoxin units (EU), a measure that does not capture LPS characteristics that influence receptor activation and possible consequences on metabolism such as changes in blood glucose.

Certain bacterial cell wall components can act as postbiotics that improve insulin sensitivity in obese mice or mice particularly when co-administered with hexa-acylated LPS (Cavallari et al., 2017). It is well established that different components of bacteria work through different host immune sensors to synergize (or tolerize) host inflammatory and metabolic responses (Cavallari et al., 2020; Schertzer et al., 2011). However, it was not known if different types of LPS interact with each other to alter blood glucose control. It is important to know if the characteristics of LPS found in different bacteria dictate metabolic and inflammatory host responses because deleterious metabolic endotoxemia is a widely used mechanism of action linking the gut microbiota and host pathology. Here, we show that the type of LPS found in different types of bacteria dictates whether metabolic endotoxemia is beneficial or deleterious since LPS characteristics influenced gut permeability, intestinal glucose absorption, blood glucose, insulin and incretins. We found that penta-acylated LPS derived from *Rhodobacter sphaeroides* acted as an insulin sensitizing postbiotic to counteract acute dysglycemia caused by hexa-acylated LPS. Endotoxemia caused by acute and chronic delivery of LPS derived from *R. sphaeroides* was metabolically beneficial and mitigated insulin resistance in obese mice.

## Results

### The type of LPS dictates changes in blood glucose, insulin, GLP-1 and adipose tissue inflammation

The acylation pattern of LPS lipid A is known to define its immune activation capacity (Lu et al., 2008; Steimle et al., 2016). We compared the impact on blood glucose regulation of the prototypical LPS from *E. coli* (with hexa-acylated lipid A) and LPS from *R. sphaeroides* (with penta-acylated lipid A) (Fig 1a). We used this comparison because the LPS from *E. coli* is widely used to model, and assumed to mediate the consequences, of metabolic endotoxemia. *E. coli*-derived LPS is known to stimulate Glucagon-like peptide (GLP)-1 secretion, elevate circulating insulin and lower blood glucose (Hagar et al., 2017; Lebrun et al., 2017; Nguyen et al., 2014). Consistently, in weight-matched mice (Fig 1b), we found that acute injection with *E. coli* LPS (0.4 mg/kg) lowered blood glucose in the fasted state and during a glucose tolerance test (GTT) compared to saline-injected mice (Fig 1c-e). Acute injection with the equivalent dose of LPS from *R. sphaeroides* did not lower fasting blood glucose and did not lower blood glucose during a GTT to the same extent compared to *E. coli* LPS during a GTT (Fig 1c-e).

**Figure 1:**
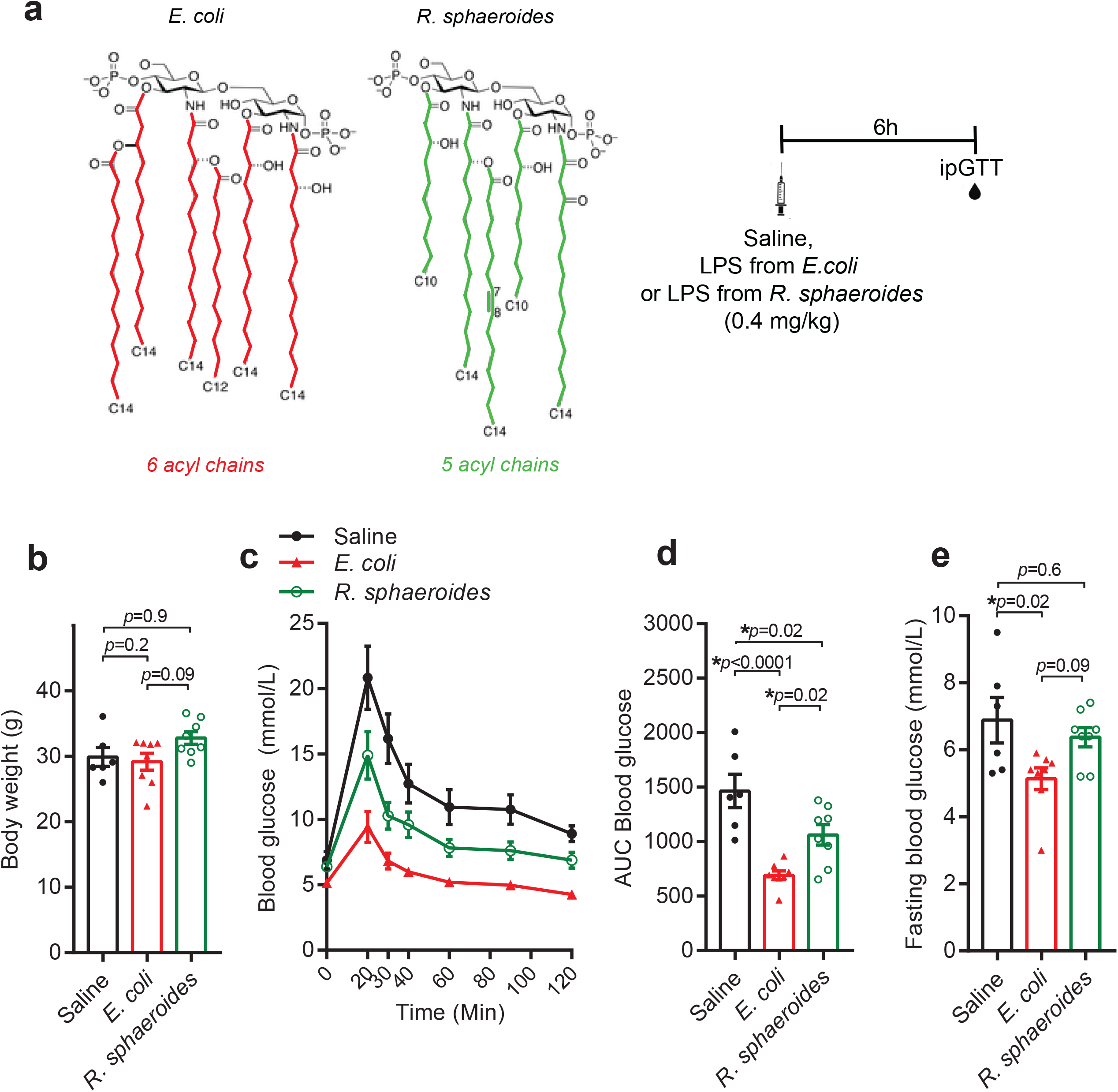
The type of LPS dictates the effect of endotoxin on blood glucose. (a) Mice were intraperitoneally injected with 0.4 mg/kg of the potent hexa-acylated LPS from *E. coli* or the under-acylated LPS from *R. sphaeroides*. Saline was injected in control mice. Six hours later, intraperitoneal glucose tolerance tests (ipGTT) were performed using 2 g/kg of glucose. (b) Body mass, (c) blood glucose and (d) area under curve (AUC) during the ipGTT. (e) Fasting blood glucose. One-way ANOVA with Tukey’s post hoc test was used to assign statistical significance, which was accepted at \**p* < 0.05. Data are shown as the mean ±SEM, and its dispersion is represented by black solid circles (saline, n=6), red solid triangles (*E. coli* LPS, n=8) or green open circles (*R. sphaeroides* LPS, n=8), where each symbol is an independent biological replicate.

We next injected a different cohort of mice with identical doses of *E. coli* LPS versus *R. sphaeroides* LPS to assess glucose-stimulated GLP-1 and insulin secretion (Fig 2a). In mice with comparable body weights (Fig 2b), *E. coli* LPS lowered blood glucose 15 min after an oral glucose load, an effect that was less pronounced using *R. sphaeroides* LPS (Fig 2c). Critically, only the LPS from *E. coli*, but not that from *R. sphaeroides*, caused an increase in GLP-1 secretion before and 15 minutes post glucose challenge in comparison with saline injected mice (Fig 2d). LPS from *E. coli* lowered blood insulin after the oral glucose challenge compared to *R. sphaeroides* LPS and saline treated mice (Fig 2e), which is likely dictated by profound lowering of blood glucose levels following *E. coli* LPS injection. In fact, blood insulin levels in saline and *R. sphaeroides* LPS injected mice were only ∼1.5-fold higher than in mice that received *E. coli* LPS (Fig 2e), but blood glucose was ∼6-fold higher (Fig 2c). Our findings corroborate the insulinogenic effect of LPS from *E. coli* (Nguyen et al., 2014) and show that the LPS derived from *R. sphaeroides* has a much lower insulinogenic potential. In line with a bacterial strain specific immunometabolic effect of different types of LPS, we found higher mRNA expression of *Il6, Il10, Nos2, Ifng, Ccl2, Cxcl1, Cxcl9* and *Cxcl10* in the visceral adipose tissue of mice after acute injection with *E. coli* LPS (0.4 mg/kg), compared to *R. sphaeroides* LPS at an equivalent dose (Fig 2f). Altogether, these data indicate that LPS derived from different bacteria and with unique acylation patterns of lipid A have divergent effects on blood glucose, GLP-1, insulin, and adipose tissue inflammation.

**Figure 2:**
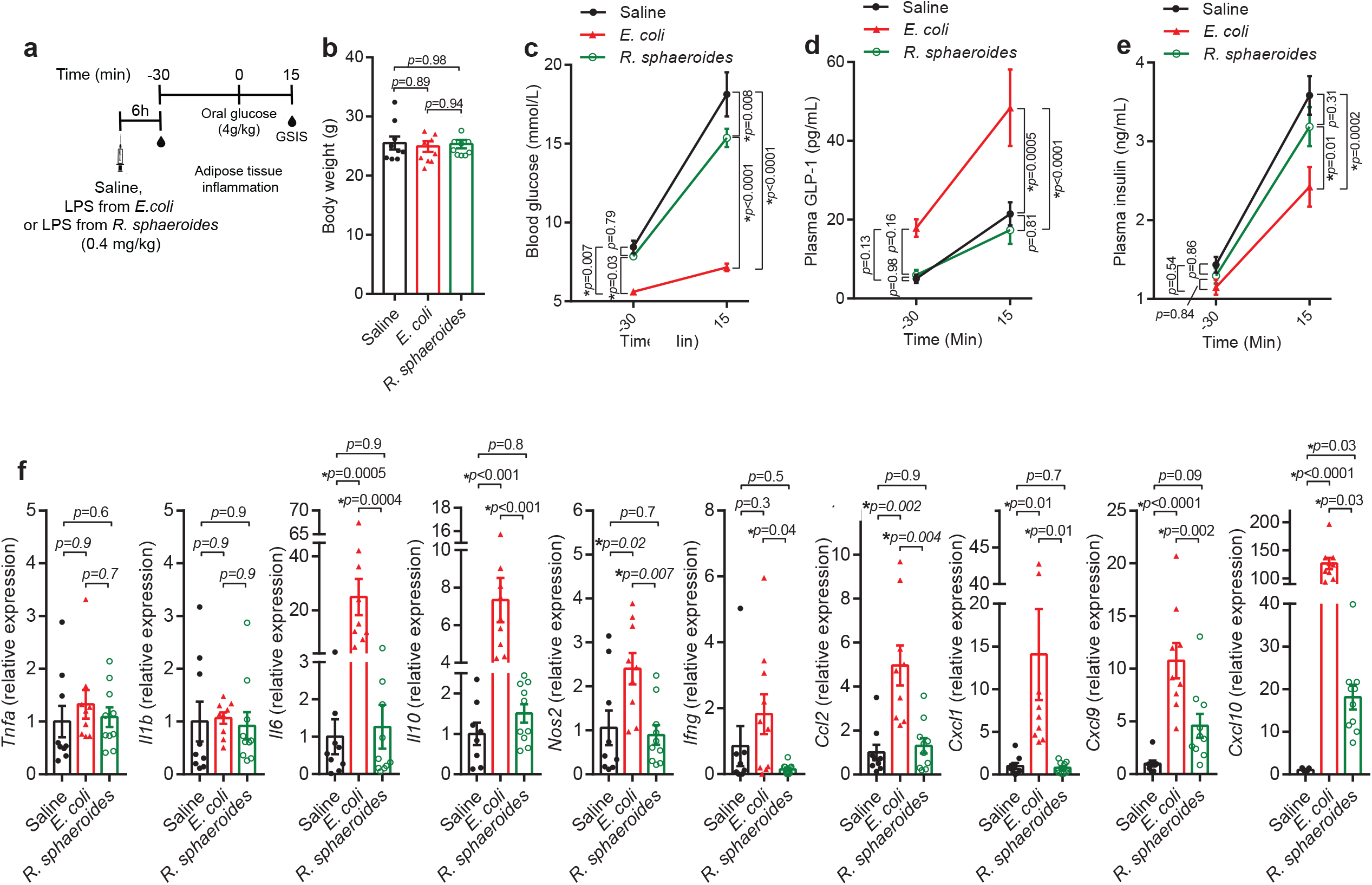
The type of LPS dictates the effect of endotoxin on insulin, GLP-1 and adipose inflammation. (a) Mice were injected with saline or acutely challenged with LPS from *E. coli* or *R. sphaeroides* (0.4 mg/kg) and basal measurements were at 30 minutes before oral glucose load (timepoint = -30 min). Glucose stimulated insulin/incretin secretion (GSIS) was assessed in blood samples collected 15 minutes after the oral glucose load (4 g/kg). (b) Body mass, (c) blood glucose (d) plasma GLP-1, and (e) plasma insulin during the GSIS. (f) Mice were injected with saline or acutely challenged with LPS from *E. coli* or *R. sphaeroides* (0.4 mg/kg), and 6 hours later gonadal adipose tissue mRNA was isolated for gene expression. Gene expression is reported relative to saline injected control mice. (b) One-way ANOVA with Tukey’s post hoc test, (c-e) two-way repeated measures ANOVA followed by Tukey’s post hoc test or (f) Kruskal-Wallis test followed by Dunn’s multiple comparisons test were used to assign statistical significance, which was accepted at \**p* < 0.05. Data are shown as the mean ±SEM, and its dispersion is represented by black solid circles (saline, n=10), red solid triangles (*E. coli* LPS, n=10) and green open circles (*R. sphaeroides* LPS, n=10), where each symbol is an independent biological replicate.

### The type of LPS dictates changes in gut barrier function

LPS derived from *E. coli* is detrimental to the gut barrier (Nighot et al., 2017), which is a key feature of metabolic endotoxemia. In weight-matched mice chronically injected with *E. coli* LPS (1 mg/kg/day for 4 days) (Fig 3a, b), we found a profound increase in paracellular intestinal permeability, as shown by higher blood plasma FITC dextran levels after an oral gavage compared to mice injected with saline (Fig 3c). Conversely, injection of an equivalent dose of *R. sphaeroides* LPS did not alter blood plasma FITC compared to saline injected mice (Fig 3c). Bacteria produce the majority of circulating D-lactate, and elevated blood D-lactate is a surrogate marker of increased intestinal permeability (Liu et al., 2016; Zhang et al., 2020). We found that only *E. coli* LPS, but not LPS from *R. sphaeroides*, increased levels of blood plasma D-lactate (Fig 3d). Taken together, this evidence supports the concept that LPS-induced gut barrier dysfunction is dependent upon the LPS type found in different bacteria.

**Figure 3:**
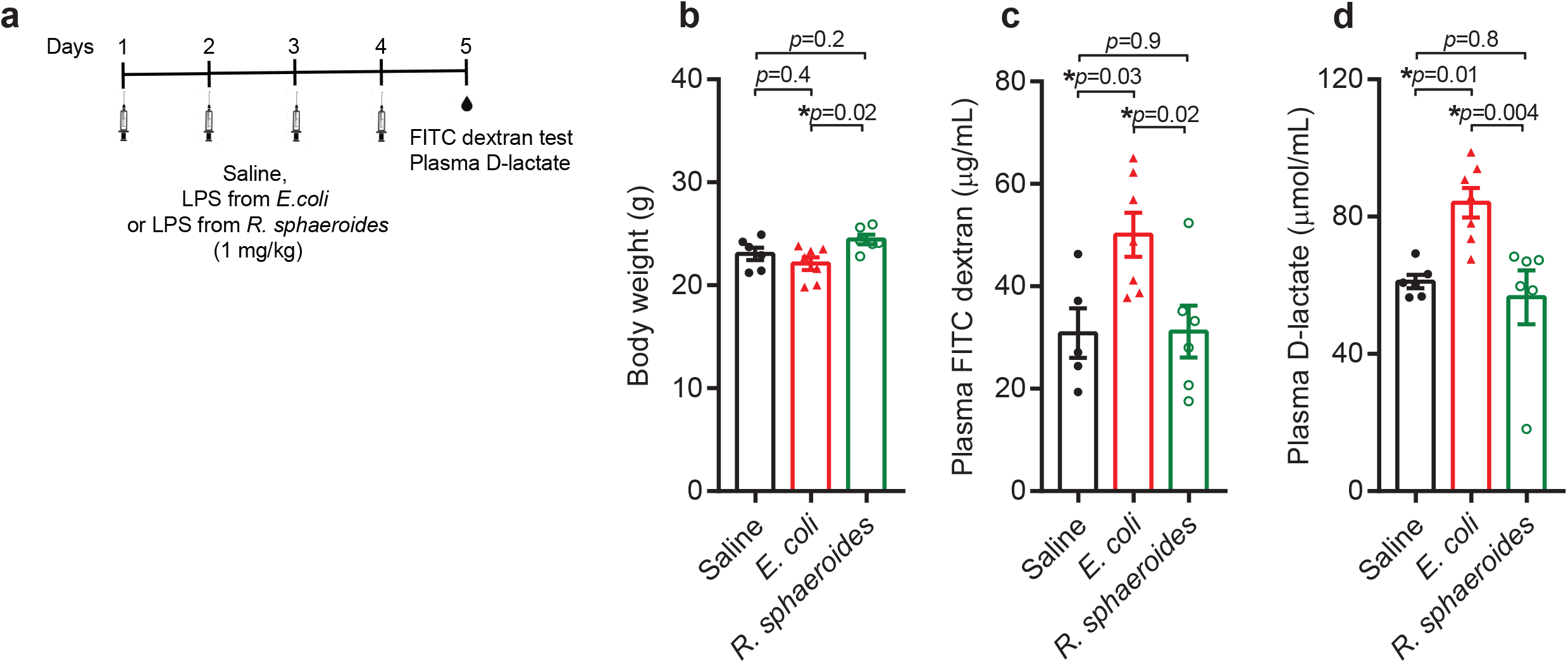
The type of LPS dictates the effect of endotoxin on gut barrier function. (a) Mice were injected with LPS from *E. coli* or *R. sphaeroides* (1 mg/kg) for four consecutive days. Control mice received saline injections. At day 5, 12h fasted mice were gavaged with FITC dextran (150 uL/mouse of a solution at 80 mg/mL) to assess intestinal permeability. Blood samples were collected before and 1h after oral FITC. Baseline relative fluorescence units were used for normalization and subtracted from FITC readings obtained 1 h after gavage. Baseline samples were also used to assess blood D-lactate. (b) Body weight during (c) FITC test. (d) Circulating D-lactate. One-way ANOVA with Tukey’s post hoc test was used to assign statistical significance, which was accepted at \**p* < 0.05. Data are shown as the mean ±SEM, and its dispersion is represented by black solid circles (saline, n=5), red solid triangles (*E. coli* LPS, n=7) and green open circles (*R. sphaeroides* LPS, n=6), where each symbol is an independent biological replicate.

### Different types of LPS interact with other bacterial components to dictate profound changes in blood glucose control

Various microbial components can interact and elicit immune synergy or tolerance to alter endocrine and metabolic responses. We have previously demonstrated that injection with the bacterial cell wall component peptidoglycan (PGN) augments blood glucose responses when combined with LPS from *E. coli* (Cavallari et al., 2017). In agreement, pre-injection with a synthetic form of PGN (FK565) caused a profound increase in blood glucose levels during a GTT following acute administration of LPS from *E. coli* (0.4 mg/kg), an effect not observed in the absence of pre-treatment with PGN (Fig 4a-d).

**Figure 4:**
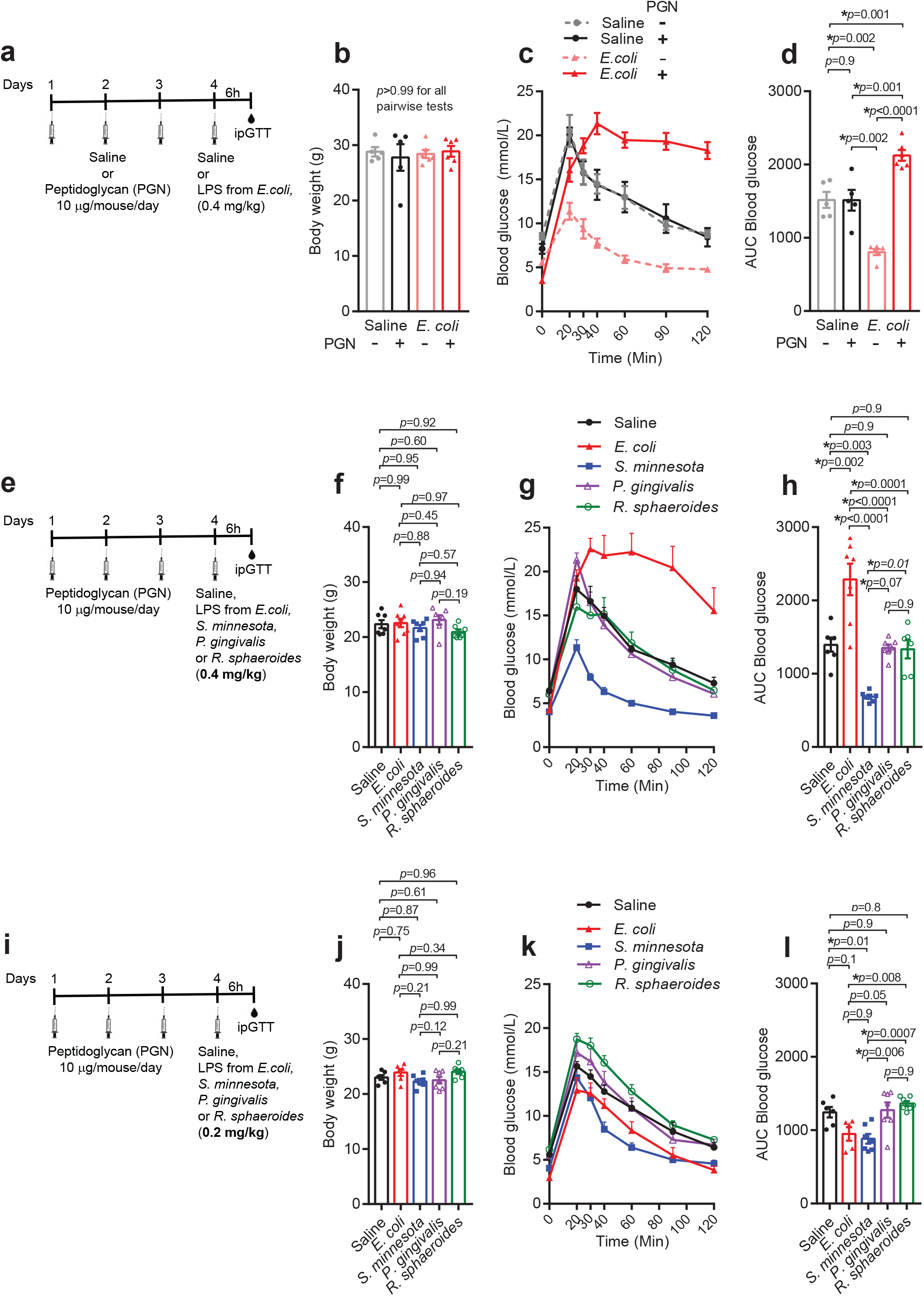
The type of LPS dictates whether synergy with other bacterial components alters blood glucose. (a) Mice were intraperitoneally injected with a synthetic peptidoglycan (PGN) that is a NOD1 agonist (FK565, 10 µg/day/mouse) or saline for three consecutive days and thereafter acutely challenged with 0.4 mg/kg of LPS from *E. coli*. Intraperitoneal glucose tolerance tests (ipGTT) were performed 6h later. (b) Body weight, (c) blood glucose and (d) area under the curve (AUC) of blood glucose during ipGTT. (e) Mice were injected with PGN (10 µg/day/mouse) for three consecutive days and then acutely challenged with 0.4 mg/kg of LPS from *E. coli, S. minnesota, P. gingivalis, R. sphaeroides*. Control mice received saline. ipGTT was performed 6h later. (f) Body weight, (g) blood glucose and (h) AUC of blood glucose during ipGTT. (i) Mice were injected with PGN (10 µg/day/mouse) for three consecutive days and thereafter acutely challenged with 0.2 mg/kg of LPS from *E. coli, S. minnesota, P. gingivalis, R. sphaeroides*. Control mice received saline. ipGTT was performed 6h later. (j) Body weight, (k) blood glucose and (l) AUC of blood glucose during ipGTT. Two-way (b, d) or one-way (f, h, j, l) ANOVA with Tukey’s post hoc test were used to assign statistical significance, which was accepted at \**p* < 0.05. Data are shown as the mean ±SEM, and its dispersion is represented in panels (b-d) by grey solid circles (saline -PGN, n=5), black solid circles (saline +PGN, n=5), rose triangles (*E*.*coli* LPS - PGN, n=5) and red triangles (*E*.*coli* LPS +PGN, n=5). Data dispersion in panels (f-h) and (j-l) is represented by black solid circles (saline, n=7 f-h, n=6 j-l); red triangles (*E. coli* LPS, n=7 f-h, n=5 j-l), blue solid squares (*S. minnesota* LPS, n=7), purple open triangles (*P. gingivalis* LPS, n=7) and green open circles (*R. sphaeroides* LPS, n=6 f-h, n=7 j-l). Each symbol is an independent biological replicate.

We next capitalized on this model to test if LPS derived from different bacterial strains altered blood glucose control. We injected mice with LPS derived from *E. coli*, the pathobionts *Salmonella minnesota* and *Porphyromonas gingivalis*, and LPS from *R. sphaeroides*, which can act as a TLR4 antagonist (Anwar et al., 2015) (Fig. 4e). In lean, male mice with similar body weight (Fig. 4f), only *E. coli* LPS at 0.4 mg/kg caused significant glucose intolerance indicated by an increased area under the curve (AUC) during a GTT, whereas the LPS from *S. minnesota* reduced glucose intolerance and the LPS from *P. gingivalis* and *R. sphaeroides* did not alter blood glucose as compared to saline controls (Figs. 4g, h). We next tested a lower dose of LPS from all the different bacterial strains (Fig 4i). In mice with similar body mass (Fig. 4j), only injection of 0.2 mg/kg LPS from *S. minnesota* reduced blood glucose after glucose load (Fig 4k, l). These results show that different types of LPS, in synergy with PGN, can affect blood glucose control. The results showing that LPS from *S. minnesota* can lower blood glucose even in the presence of PGN are interesting. LPS from *S. minnesota* R959 contains a mixture of hepta-acyl and hexa-acyl lipid A components (Qureshi et al., 1985), which makes it a complex system to test. We therefore opted to further test the LPS derived from *R. sphaeroides*, which contains penta-acylated lipid A.

### Different types of LPS interact with each other to dictate changes in blood glucose control

We next assessed the potential for synergy or antagonism between different types of LPS on glucose control in mice pre-injected with PGN (Fig. 5a). As expected, injecting mice with 0.4 mg/kg of LPS derived from *E. coli* caused profound glucose intolerance and increased the AUC during a GTT, but injection of *R. sphaeroides* LPS did not alter blood glucose in mice with comparable body mass (Fig. 5b-d). Further, *R. sphaeroides* LPS could counteract the dysglycemia caused by *E. coli* LPS. Despite mice having twice the dose of LPS, co-injection with 0.4 mg/kg each of *E. coli* and *R. sphaeroides* actually mitigated higher blood glucose during a GTT such that the combination of these two types of LPS did not significantly alter AUC compared to saline injection (Figs. 5b-d). At a lower dose of each type of LPS (0.2 mg/kg, Fig 5e), co-injection of both *E. coli* LPS and *R. sphaeroides* LPS caused a significant reduction in blood glucose during a GTT compared to mice injected with saline and with comparable body mass (Fig 5f-h). Importantly, co-injection with *R. sphaeroides* LPS and *E. coli* LPS (each at 0.2 mg/kg) actually lowered blood glucose during a GTT despite the total dose of LPS (0.4 mg/kg), which would cause profound dysglycemia if it were all derived from *E. coli* (Fig 5f-h). An injection of 0.2 mg/kg of *R. sphaeroides* LPS alone still had no effect on blood glucose or AUC during a GTT in mice (Figs. 5f-h). This data shows that different types of LPS can antagonize each other to influence blood glucose, particularly at lower doses of LPS that mimic metabolic endotoxemia.

**Figure 5:**
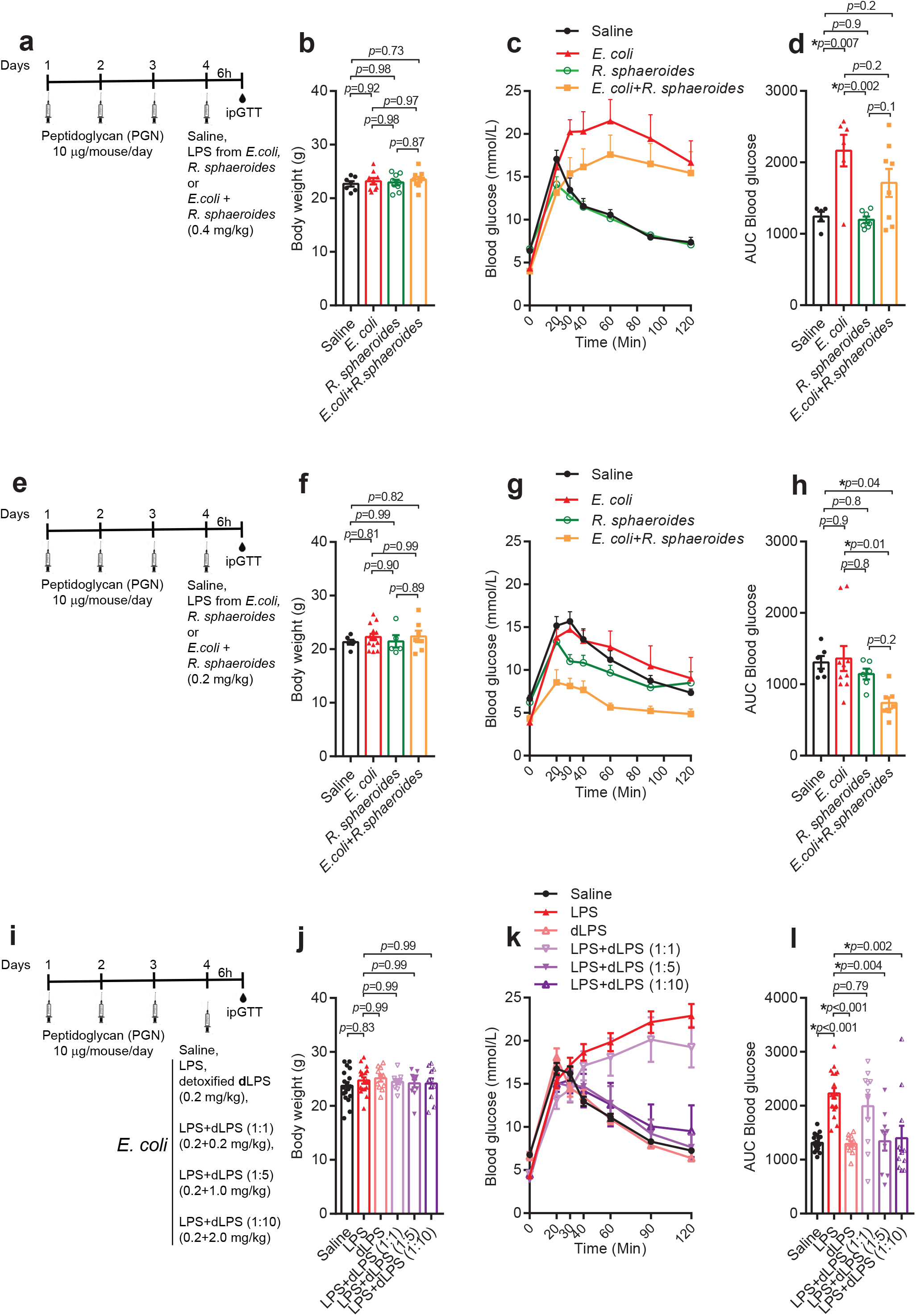
Different types of LPS interact to dictate changes in blood glucose control. Mice were intraperitoneally injected with a synthetic peptidoglycan (PGN) that is a NOD1 agonist (FK565, 10 µg/day/mouse) for three consecutive days and thereafter acutely challenged with (a) 0.4 or (e) 0.2 mg/kg of LPS derived from *E. coli, R. sphaeroides* or a combination of both types of LPS at equivalent doses. Control mice were injected with saline. Intraperitoneal glucose tolerance tests (ipGTT) were performed 6h later. (b, f) Body mass, (c, g) blood glucose, (d, h) area under the curve (AUC) of blood glucose. (i) Mice pre-treated with PGN were injected with hexa-acylated or detoxified (*i*.*e*., partially delipidated) LPS from *E. coli* (dLPS) at 0.2 mg/kg or a combination of LPS+dLPS at a proportion of 1:1, 1:5 or 1:10. Control mice were injected with saline. IpGTT were performed 6h later. (j) Body mass, (k) blood glucose, (l) AUC of blood glucose. (b, f, j) One-way ANOVA with Tukey’s post hoc test or (d, h, l) Kruskal-Wallis followed by Dunn’s multiple comparisons test were used to assign statistical significance, which was accepted at \**p* < 0.05. Data are shown as the mean ±SEM, and its dispersion is represented by black solid circles (saline, n=6 b-h, n=18 j-l), red solid triangles (*E. coli* LPS, n=10 b-d, n=6 f-h, n=18 j-l), green open circles (*R. sphaeroides* LPS, n=6 b-d, n=8 f-h), solid orange squares (*E. coli* + *R. sphaeroides* LPS, n=9), open rose triangles (dLPS, n=11), open light purple inversed triangles (LPS+dLPS 1:1, n=11), solid purple inversed triangles (LPS+dLPS, n=11) and half solid/half open dark purple triangles (LPS+dLPS 1:10, n=11) where each symbol is an independent biological replicate.

Since lipid A deacylation is a well-known host strategy to detoxify LPS, we sought to test the impact of the interaction between hexa-acylated and detoxified (*i*.*e*., partially delipidated) LPS from *E. coli* on blood glucose of weight-matched mice pre-injected with PGN (Fig 5i). We used a commercially available detoxified *E. coli* LPS (dLPS), where the lipid A moiety can harbor 1-5 acyl chains (Sigma-Aldrich, Cat# L3023). As expected, injection of hexa-acylated 0.2 mg/kg *E. coli* LPS in weight-matched mice caused profound glucose intolerance, whereas an equivalent dose of dLPS had no impact on blood glucose during a GTT (Figs. 5j-k). We found that while co-injection with 0.2 mg/kg of *E*.*coli* LPS+dLPS at 1:1 ratio still caused profound glucose intolerance, but injection of *E. coli* LPS together with dLPS at a 1:5 or 1:10 ratio prevented *E. coli* LPS-induced glucose intolerance during a GTT (Figs. 5j-k). These findings show that the ratio of different types of LPS, rather than simply the total LPS dose, can determine the impact of endotoxins on glycemia, particularly at lower doses of LPS, which more closely resemble metabolic endotoxemia. Our findings also show that LPS deacylation contributes to impact of LPS from the same type of bacteria on blood glucose. Therefore, host mechanisms of LPS detoxification such as deacylation are positioned to influence how metabolic endotoxemia alters blood glucose.

### Endotoxin units do not capture the potential of different LPS types to alter blood glucose

We next assessed the supposed potency of LPS from *E. coli* and *R. sphaeroides* using a Limulus amebocyte lysate (LAL)-based assay to quantify Endotoxin Units (EU). The under-acylated (i.e. penta-acylated) LPS derived from *R. sphaeroides* produced ∼1×10^6^ EU per mL, which was about half as potent than hexa-acylated LPS from *E. coli* (Fig 6a). Importantly, *R. sphaeroides* LPS still produced an EU value despite our results showing no effect on blood glucose and despite the known antagonistic action of *R. sphaeroides* LPS on TLR4 (Anwar et al., 2015). Since LPS from *E. coli* and *R. sphaeroides* have distinct lipid A acylation patterns (Fig. 1a), we next matched the potency of these two LPS molecules based on the EU values obtained. We calculated that 0.9×10^6^ EU equates to 0.4 and 0.8 mg of LPS from *E. coli* and *R. sphaeroides*, respectively. Injection of 0.9×10^6^ EU of either type of LPS in mice pre-treated with PGN (Fig 6b) did not alter body mass (Fig. 6c), but only *E. coli* derived LPS caused glucose intolerance, whereas LPS from *R. sphaeroides* did not alter blood glucose or AUC during a GTT in lean mice (Figs. 6d, f). These results show that the effect of LPS on blood glucose regulation is dependent on the bacterial source, and that EU quantification does not reflect how LPS or metabolic endotoxemia from different bacterial strains alter blood glucose control.

**Figure 6.**
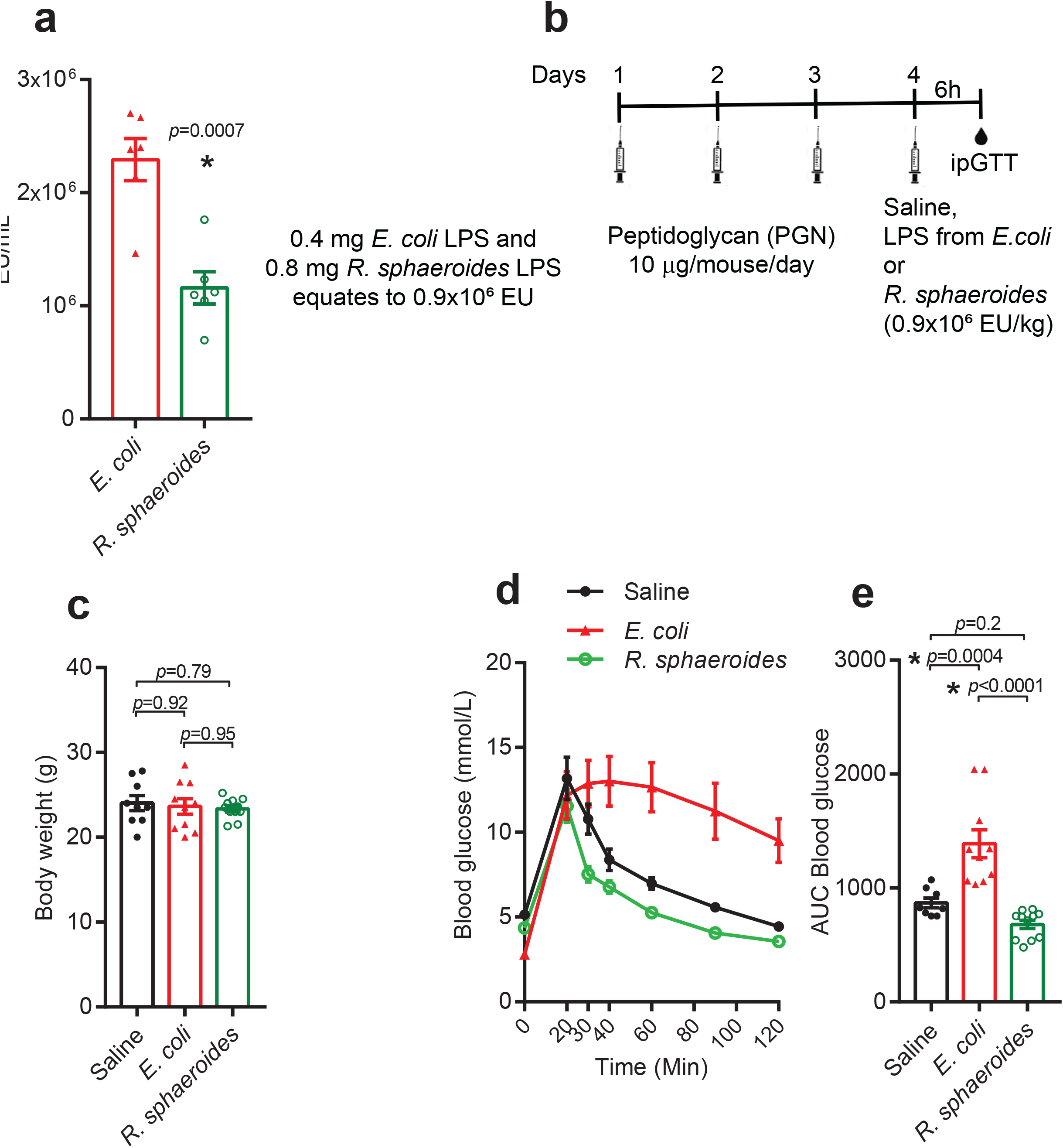
Endotoxin units do not capture the potential of different types of LPS to modify blood glucose. (a) Quantification of endotoxin units (EU)/mL in 1 mg/mL solutions of LPS from *E. coli* and *R. sphaeroides* (n=5, technical replicates). (b) Mice were intraperitoneally injected with a synthetic peptidoglycan (PGN) that is a NOD1 agonist (FK565, 10 µg/day/mouse) for three consecutive days and thereafter acutely challenged with 0.9×10^6^ EU/mL of LPS from *E. coli* or *R. sphaeroides*. This dose equates to 0.4 and 0.8 mg/kg of *E. coli* and *R. sphaeroides* LPS, respectively. (c) Body mass, (d) blood glucose, (e) area under the curve (AUC) of blood glucose. (a) Mann-Whitney U test, (c) One-way ANOVA with Tukey’s post hoc test or (e) Kruskal-Wallis followed by Dunn’s multiple comparisons test were used to assign statistical significance, which was accepted at \**p* < 0.05. Data are shown as the mean ±SEM, and its dispersion is represented by black solid circles (saline, n=9), red solid triangles (*E. coli* LPS, n=10), and green open circles (*R. sphaeroides* LPS, n=10), where each symbol is an independent biological replicate.

### The type of LPS dictates their impact on intestinal glucose absorption

We showed that gut permeability and blood glucose during an oral glucose load were influenced by the type LPS. Hence, we sought to determine the impact of LPS on intestinal glucose absorption. Mice were acutely injected with an EU-matched dose of *E. coli* or *R. sphaeroides* LPS and, 6h later, received an oral dose of a non-metabolizable glucose analog (3-o-methyl-d-glucopyranose [3-OMG]) plus D-glucose (4 g/kg); 3-OMG appearance in the blood was monitored to indicate intestinal glucose absorption (Fig 7a). In weight-matched mice, we found lower blood glucose levels during the GTT in *E. coli* LPS treated mice compared to saline and *R. sphaeroides* LPS injected mice (Fig 7 b-d). At this dose, *R. sphaeroides* LPS still lowered blood glucose during the GTT compared with saline treated mice, but to a lesser extent than *E. coli* LPS treated mice (Fig 7b-d). These findings are in line with the glucose tolerizing effect seen after 0.4 mg/kg of either type of LPS (Fig 1). In saline- and *R. sphaeroides* treated mice, serum levels of 3-OMG peaked 40 min after oral delivery of the glucose analog, whereas a delayed peak (≥ 90 min) was observed in *E. coli* LPS treated mice (Fig 7e). The rate of appearance of 3-OMG in the blood was approximately two times lower in *E. coli* treated mice as compared with saline and *R. sphaeroides* injected mice (Fig 7f). An EU-matched dose of *R sphaeroides* LPS did not alter intestinal glucose absorption as compared with saline injected mice (Fig 7e, f). This data shows that delayed intestinal glucose absorption may contribute to lower blood during GTT after acute *E. coli* LPS treatment. However, *R. sphaeroides* LPS did not alter intestinal glucose absorption and can still lower blood glucose during a GTT. These data reinforce that *R. sphaeroides* has blood glucose lowering properties independent of changes in intestinal glucose absorption. To the best of our knowledge, this is the first *in vivo* demonstration that the type of LPS dictates changes in intestinal glucose absorption.

**Figure 7:**
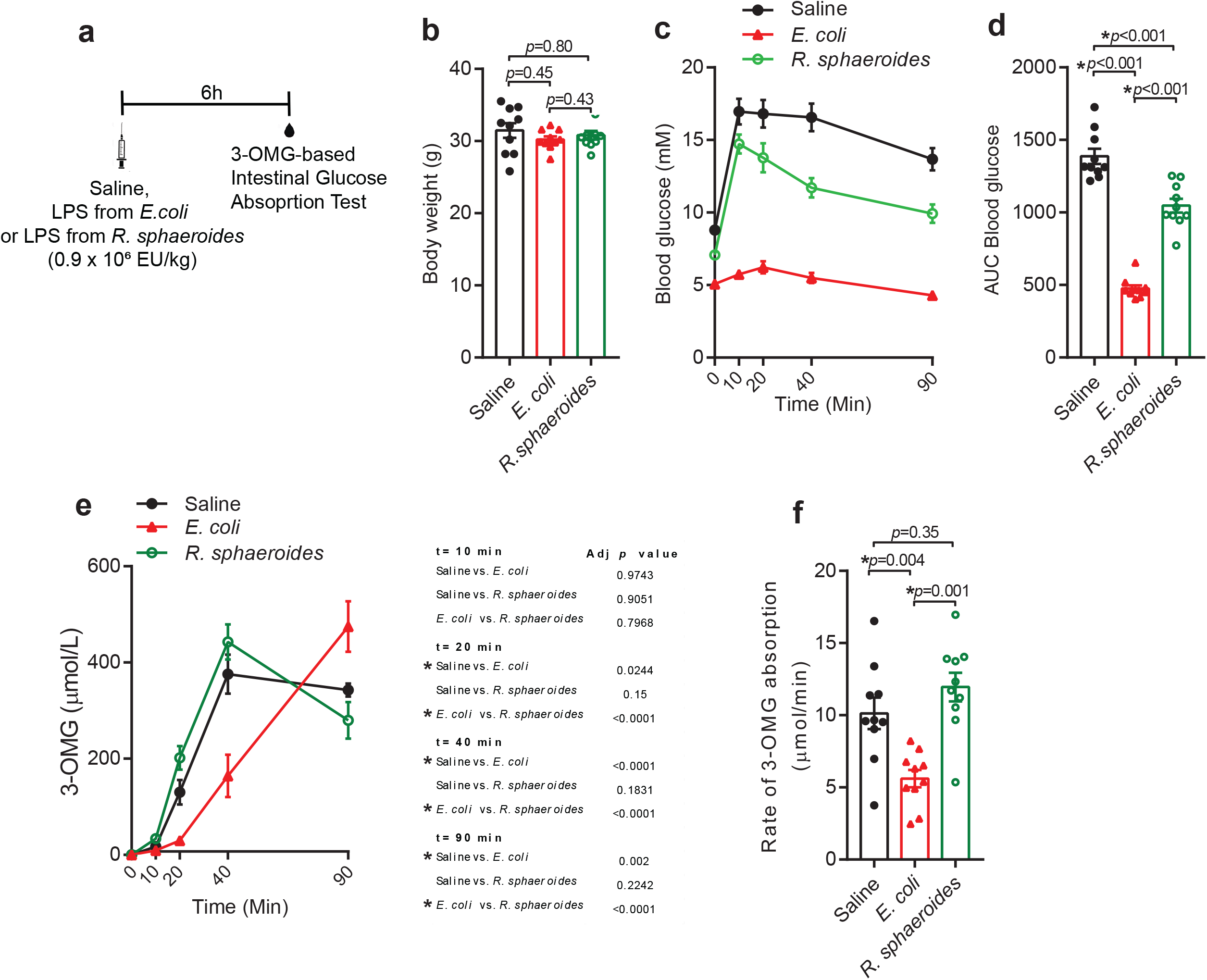
LPS type alters intestinal glucose absorption. (a) Lean mice fed a standard chow diet were acutely injected with 0.9×10^6^ EU/kg of *E. coli* or *R. sphaeroides* LPS, or saline. After 6h, mice received a gavage of 4 mg/mouse of 3-o-methyl-d-glucopyranose (3-OMG) and 4 g/kg d-glucose. (b) Body mass, (c) blood glucose, and (d) area under the curve (AUC) of blood glucose during the 3-OMG containing glucose tolerance test (GTT). (e) 3-OMG levels in the blood. (f) Rate of 3-OMG appearance in the blood calculated from time 0 to the peak value of 3-OMG. *P* values were calculated using (b-d, f) one-way ANOVA with Tukey’s post hoc test or (e) two-way repeated measures ANOVA with Tukey’s multiple comparisons test, and significance was accepted at \**p* < 0.05. Data are shown as the mean ±SEM, and its dispersion is represented by black solid circles (saline, n=10), red solid triangles (*E. coli* LPS, n=10), and green open circles (*R. sphaeroides* LPS, n=10), where each symbol is an independent biological replicate.

### Metabolically beneficial endotoxemia caused by *R. sphaeroides* LPS improves insulin sensitivity in obese mice

We showed that LPS from *R. sphaeroides* can antagonize dysglycemia caused by *E. coli* LPS in lean mice. Therefore, we next assessed whether the LPS from *R. sphaeroides* could be used as an insulin-sensitizing postbiotic treatment when administered acutely to diet-induced obese mice (Fig 8a). We matched the doses of *E. coli* and *R. sphaeroides* LPS by EU (0.9×10^6^ EU/kg). Compared to saline-treated mice, injection of *R. sphaeroides* LPS lowered blood glucose during an oral glucose load, whereas *E. coli* derived LPS lowered blood glucose to an even greater extent, but neither type of LPS altered body mass in obese mice (Fig 8b-d). Furthermore, only injection of *E. coli* derived LPS increased oral glucose-stimulated plasma insulin compared with saline-treated obese mice, an effect not seen in obese mice injected with LPS from *R. sphaeroides* (Fig. 8e, f). Injection of LPS from *R. sphaeroides* or *E. coli* did not alter HOMA-IR, assessed in fasted obese mice (Fig. 8g). However, *E. coli* derived LPS increased the insulinogenic index and insulin resistance index during the oral glucose load (Fig. 8h,i). In contrast, LPS from *R. sphaeroides* did not alter the insulinogenic index and lowered the insulin resistance index during the oral glucose load in obese mice (Fig. 8h, i). These findings indicate that the LPS from *R. sphaeroides* can mitigate obesity-driven insulin resistance and lower blood glucose without increasing blood insulin during a glucose load. We next injected a different set of obese mice, fed an obesogenic diet for a longer period of time, with *E. coli* or *R. sphaeroides* LPS (0.9×10^6^ EU/kg) and assessed insulin signaling in the visceral adipose tissue upon insulin stimulation (Fig 8j). In this cohort of weight-matched extremely obese mice (Fig 8k), acute injection with *E. coli* LPS did not alter insulin-stimulated phosphorylation of AKT^ser473^, but acute administration with a single EU-matched dose of *R. sphaeroides* LPS significantly increased insulin-stimulated phosphorylation of AKT^ser473^ in adipose tissue compared to mice injected with saline or *E*.*coli* LPS (Fig 8l). These findings show that *R. sphaeroides* LPS can increase adipose tissue insulin signaling during protracted diet-induced obesity in mice.

**Figure 8:**
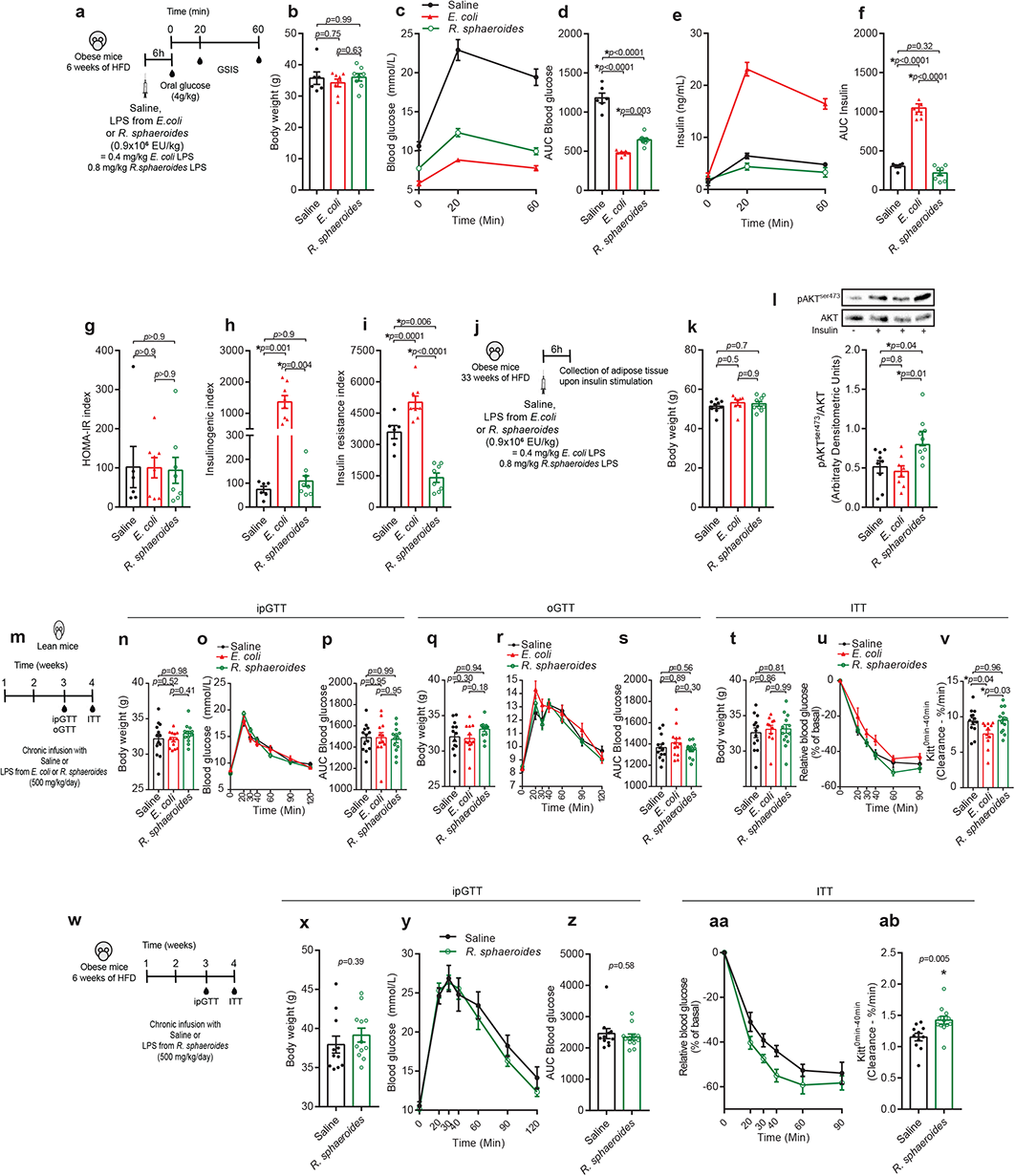
Metabolically beneficial endotoxemia caused by *R. sphaeroides* LPS improves insulin sensitivity in obese mice. (a) Diet-induced obese mice were fed a high fat det (HFD) for 6 weeks and acutely challenged with saline, *E. coli* or *R. sphaeroides* LPS (0.9×10^6^ EU/kg). Glucose-stimulated insulin secretion (GSIS) was performed by assessing plasma insulin before (0 minutes) and after (20, 60 minutes) an oral glucose load (4 g/kg). (b) Body mass, (c) blood glucose, (d) area under blood glucose curve (AUC), (e) plasma insulin and (f) AUC of insulin during the GSIS conducted 6 h after LPS injection. (g) HOMA-IR (fasting insulin x fasting glucose/22.5). (h) Insulinogenic index (ΔInsulin/ΔGlucose) and (i) insulin resistance index (AUC insulin x AUC glucose/100) during the GSIS. (j) Gonadal adipose tissue was collected after acute insulin injection in obese mice and 6h after injection with saline or LPS from *E. coli* or *R. sphaeroides* (0.9×10^6^EU/kg). (k) Body weight and (l) pAKT^ser473^ and total AKT levels were assessed in adipose tissue. (m) Osmotic mini pumps were subcutaneously inserted in lean mice that received 500 µg/kg/day of *E. coli* LPS, *R. sphaeroides* LPS or saline for 4 weeks. (n) Body weight, (o) blood glucose levels (p) AUC during intraperitoneal (ip)GTT. (q) Body weight, (r) blood glucose levels (s) AUC during oral (o)GTT. (t) Body weight, (u) relative change in blood glucose and (v) glucose disappearance rate (Kitt) during insulin tolerance test (ITT). (w) Osmotic mini pumps were subcutaneously inserted in obese mice that were fed a HFD for 6 weeks and received 500 µg/kg/day of *R. sphaeroides* LPS or saline for 4 weeks. (x) Body mass, (y) blood glucose (z) and AUC of blood glucose during ipGTT. (aa) Relative change in blood glucose and (ab) glucose disappearance rate during an ITT. (b, k, n, q, p, s, t, v) One-way ANOVA with Tukey’s post hoc test, (d, f, g-i) Kruskal-Wallis test followed by Dunn’s multiple comparisons test or (x, z, ab) unpaired parametric t-test were used to compare groups. Statistical significance was accepted at \**p* < 0.05. Data are shown as the mean ±SEM, and its dispersion is represented by black solid circles (saline, n=6 b-i, n=9 k, l, n=10 n-v n=11 x-ab), red solid triangles (*E. coli* LPS, n=8 b-i, n=8 k, l, n=10 n-v) and green open circles (*R. sphaeroides* LPS, n=8 b-i, n=9 k, l, n=10 n-v, n=11 x-ab), where each symbol is an independent biological replicate.

It is known that chronic subcutaneous administration of a low dose of *E. coli* LPS models metabolic endotoxemia and can promote insulin resistance in lean mice (Cani et al., 2007). We next tested if chronic subcutaneous infusion of *R. sphaeroides* LPS (500 µg/kg/day) could alter insulin resistance in lean mice (Fig 8m). To gain insight into LPS-induced changes in glycemic control, we assessed glucose tolerance in lean mice using intraperitoneal (Fig 8 n-p) or the oral (Fig 8 q-s) routes to deliver glucose. We found that chronic infusion with *E. coli* or *R. sphaeroides* LPS did not alter blood glucose levels during an intraperitoneal or oral GTT in weight-matched mice (Fig 8n-s). Weight-matched, lean mice chronically treated with *E*.*coli* LPS had lower blood glucose disappearance during an ITT compared to mice treated with *R. sphaeroides* LPS or saline (Fig 8t-v). Further, *R. sphaeroides* LPS did not alter glucose clearance during ITT as compared to saline treated mice (Fig 8t-v). These data indicate that chronic treatment with *E. coli* LPS impairs insulin tolerance in lean mice, but *R. sphaeroides* LPS did not alter glycemia or insulin tolerance in lean mice. Finally, we investigated chronic infusion of *R. sphaeroides* LPS during diet-induced obesity in mice (Fig. 8w). Infusion of LPS from *R. sphaeroides* for 4 weeks did not alter body mass or glucose tolerance in mice fed an obesogenic high fat diet for a total of 10 weeks (Fig 8w-z). However, chronic delivery of LPS from *R. sphaeroides* increases blood glucose disappearance rate during an ITT compared to saline-infused mice (Fig 8aa, ab). These data show that metabolically beneficial endotoxemia by chronic elevation of LPS from certain types of bacteria (such as *R. sphaeroides*) is sufficient to improve insulin tolerance in obese mice.

## Discussion

The concept of metabolic endotoxemia is often portrayed as deleterious actions of gut microbiota-derived LPS, which promotes inflammation, dysglycemia and metabolic dysfunction. We propose that the concept of metabolic endotoxemia should include LPS characteristics that discriminate the deleterious versus beneficial features of LPS and endotoxemia. Our results show that the type of LPS found in different bacteria regulates changes in gut barrier function, inflammation, hormones, and blood glucose. Hence, the concept of metabolic endotoxemia should move beyond LPS load to include LPS characteristics, where the net effect on metabolism can be influenced by a complex and dynamic gut microbiota. Our results also have implications for the modelling and measurement of metabolic endotoxemia, which is often reported in terms of EU that is presumed to reflect the activity of total LPS levels in circulation. Here we show that certain types of LPS can be metabolically beneficial despite measurable EU and we show that LPS from different types of bacteria matched for EU have distinct effects on gut barrier function, adipose inflammation, intestinal glucose absorption, blood glucose, insulin, and incretins. Many of these LPS-type dependent hormonal, glucose, inflammation and barrier function responses are likely connected. For example, delayed intestinal glucose absorption caused by *E. coli* LPS may potentiate incretin and insulin responses.

Our findings revealed that LPS from certain bacteria can promote a metabolically beneficial endotoxemia, which can counteract metabolically detrimental endotoxemia such as *E. coli* LPS-driven dysglycemia. Hence, LPS from different types of bacteria can interact and we show this has profound consequences on blood glucose when combined with other bacterial components, including as parts of the bacteria cell wall. These interactions should be considered when assessing host metabolism, metabolic endotoxemia and dynamic changes in the gut microbiota. Obesity increases levels of other bacterial components, such as NOD1-activating peptidoglycan (Chan et al., 2017) and bacterial DNA (Anhê et al., 2020b; Denou et al., 2015) in the blood and various host tissues. The synergy and tolerance induced by the interaction between bacterial components, including different types of LPS in each tissue, should be examined, which could help refine the concept of bacterial postbiotics and metabolic endotoxemia.

We propose that defining metabolic endotoxemia by LPS characteristics is more relevant to metabolic outcomes compared to EU quantification. As one example, LPS from *R. sphaeroides* promoted insulin sensitivity in obese mice. Lipid A characteristics can vary in the number, length, saturation, and distribution of fatty acid chains, which can determine LPS immunogenicity and engagement of MD2/TLR4 (Steimle et al., 2016). In general, the immunogenicity increases as the number of phosphate groups and acyl chains increase on the lipid A moieties from different bacteria (Berezow et al., 2009). While all these structural characteristics can be targets for detoxification and influence LPS potency, our data shows that the number of lipid A acyl chains is a pivotal mediator that dictates the effects of metabolic endotoxemia on blood glucose. The host possesses a conserved enzymatic system devoted to inactivating LPS by promoting its deacylation (Lu et al., 2008). We showed that partially delipidated LPS from *E. coli* can prevent dysglycemia caused by the intact hexa-acylated LPS from the same bacterial strain, which implies that host LPS detoxification processes can tip the balance of detrimental versus beneficial endotoxemia. Our results show that specific bacteria with favorable acyl chain characteristics on lipid A (i.e., under-acylated) represent new potential targets for prebiotics or probiotics. There are several other host LPS detoxification mechanisms, including intestinal alkaline phosphatases and lipopolysaccharide binding protein (LBP) that warrant investigation (Goldberg et al., 2008; Lamping et al., 1998). Further, our results suggest that certain markers of metabolic endotoxemia should be re-evaluated given that EU or circulating LBP may be of limited use as standalone markers of metabolic endotoxemia since neither are not positioned to discriminate different types of LPS.

In our mouse models, only the LPS from *E. coli* caused dysglycemia and insulin resistance. LPS from *P. gingivalis* had no impact on blood glucose control, and the LPS derived from *R. sphaeroides* lowered insulin resistance in obese mice. These findings highlight the potential for divergence in the lipid A structure of LPS from different bacteria to reach beyond regulation of innate immunity and include effects on blood glucose. Interestingly, in healthy humans, TLR4 antagonism driven by under-acylated LPS was the dominant signal derived from the entire fecal bacterial community (d’Hennezel et al., 2017), suggesting that the net effect of gut microbiota-derived LPS is to facilitate host tolerance and promote insulin sensitivity. Hence, these data and concepts warrant careful examination of strain specific LPS characteristics, such as acylation status, rather than just LPS load (defined by EU), in the inflammatory and metabolic consequences of metabolic endotoxemia during obesity.

The demonstration that the penta-acylated and under-saturated LPS from *R. sphaeroides* can protect from dysglycemia and insulin resistance in mice provides proof-of-principle for one type of LPS that can mitigate the plethora of microbial derived molecules and other factors promoting inflammation and insulin resistance during obesity. There are several pieces of evidence linking LPS-containing Gram-negative bacteria to improved glucose control. Bariatric surgery lowers blood glucose in parallel to expanding Proteobacteria (Anhê et al., 2017). Human-to-mice transfer of fecal microbes that persist after bariatric surgery is sufficient to lower blood glucose in pre-clinical rodent models (Arora et al., 2017; de Groot et al., 2020), and the LPS characteristics of this bacterial population should be considered. Furthermore, pasteurized bacterial components from the Gram-negative bacterium *Akkermansia muciniphila* were shown to improve markers of glycemia (Depommier et al., 2019). Hence the ability of prebiotics, probiotics and postbiotics to influence LPS characteristics should be considered in metabolic endotoxemia. Our data highlights that lipid A structure, and in particular the acylation pattern, may guide the search for bioactive components in Gram-negative probiotic strains or novel probiotic bacteria with insulin sensitizing potential.

## Acknowledgements

We are thankful to Dr Patricia Brubaker, Dr Alexander Martchenko and Dr Sarah E. Wheeler for technical assistance with GLP-1 assessment.

## Funding

This work was supported by grants to J.D.S. from the Canadian Institutes of Health Research (FDN-154295). F.F.A. holds a CIHR postdoctoral fellowship. J.D.S. holds a Canada Research Chair in Metabolic Inflammation.

## Author Contributions

F.F.A. derived the hypothesis, conducted experiments, analyzed the data, contributed to the discussion, and wrote the manuscript. J.F.C. and N.G.B. helped with experiments and contributed to the discussion. J.D.S. researched the data, derived the hypothesis, and edited the manuscript.

## Competing interest

The authors declare no conflict of interest.

## Methods

### RESOURCE AVAILABITY

#### Lead contact

Further information and requests for resources and reagents should be directed to and will be fulfilled by the Lead Contact, Dr Jonathan D. Schertzer.

#### Data and code availability

The datasets generated and/or analyzed during the current study are available from the corresponding author on reasonable request. This study did not generated/analyzed codes.

### EXPERIMENTAL MODEL AND SUBJECT DETAILS

#### Animals

All procedures were approved by the McMaster University Animal Research Ethics Board (Animal Utilization Protocol #16-02-96). Nine- to sixteen-week-old C57BL/6NTac mice were purchased from Taconic and sourced from in-house colonies established with mice received from Taconic. All mice were kept on a control diet (Teklad 22/5, catalog #8640) or high fat diet (D12492, Research diets).

## METHOD DETAILS

### Acute LPS treatment in lean mice

All LPS were ultrapure grade supplied by InvivoGen (San Diego, CA), except for LPS utilized in Figure 5, panels i-l (Sigma-Aldrich). LPS potency was determined using a Limulus amebocyte lysate-based (LAL) assay (Pierce Chromogenic Endotoxin Quant Kit, Thermo Scientific). Mice were intraperitoneally (ip) injected with LPS from various bacterial strains at different doses and subjected to metabolic tests 6h post LPS challenge. In figures 1 and 2, mice received 0.4 mg/kg LPS from *Escherichia coli* O111:B4 or *Rhodobacter sphaeroides*. In Figure 7, mice received EU-matched does (0.9×10^6^ EU/kg) of these LPS.

### Acute LPS treatment in obese mice

Fourteen-week-old diet-induced obese mice (fed the obesogenic high fat diet – HFD - for 8 weeks) were transferred to clean cages without food and injected with either saline, *E. coli* or *R. sphaeroides* LPS at 0.9×10^6^ EU/kg. Six hours later, mice were subjected to glucose-stimulated insulin secretion tests. Alternatively, the same procedure was carried out in 39-week-old obese mice, fed on HFD for 33 weeks. Adipose tissue was collected and used to assess insulin signaling by immunoblotting.

### Acute LPS treatment in lean mice pre-treated with peptidoglycan

Peptidoglycan (FK565) was supplied by Astellas Pharma (Osaka, Japan) and ip administered to mice for three consecutive days at 10 µg/day/mouse. In figure 4a-d, following peptidoglycan treatment, mice were ip injected with LPS from *E. coli* or saline and, after 6h, subjected to metabolic tests. In figure 4e-l and figure 5, mice pre-treated with peptidoglycan were ip injected with LPS from *E. coli, Salmonella minnesota* R595, *Porphyromonas gingivalis* or *R. sphaeroides* at 0.2 or 0.4 mg/kg. In Figure 5 (a-h), in addition to saline, *E. coli* and *R. sphaeroides* LPS, mice were also co-injected with equivalent doses of *E. coli* and *R. sphaeroides* LPS. In figure 5 (i-l), mice received hexa-acylated or detoxified (*i*.*e*., partially delipidated) LPS from *E. coli* (dLPS) at 0.2 mg/kg or a combination of LPS+dLPS at a proportion of 1:1, 1:5 or 1:10. In figure 6, mice pre-injected with peptidoglycan received saline or LPS from *E. coli* or *R. sphaeroides* at 0.9×10^6^ EU/kg. Glucose tolerance tests (GTT) were performed 6 hours after LPS injection.

### Chronic LPS treatment

Lean mice received four consecutive injections with 1 mg/kg LPS from *E. coli* or *R. sphaeroides* followed by *in vivo* gut permeability assay at day 5. Alternatively, osmotic mini pumps (Alzet, model 2004) were loaded with saline, LPS from E. coli or LPS from *R. sphaeroides* and subcutaneously inserted into 20-week-old lean mice. Mini pumps containing saline or LPS from *R. sphaeroides* were subcutaneously inserted in 12-week-old obese mice that had already been fed high fat diet for 6 weeks. Mice were chronically exposed to saline or LPS (500 µg/kg/day) for 4 weeks via continual release by osmotic mini pumps, where mice were continually fed a standard chow diet (lean mice) or a high fat diet (obese mice).

### Glucose (GTT) and insulin (ITT) tolerance tests

Fasted mice (8:00 to 14:00 for GTT; 3:00 to 9:00 for ITT) were injected with glucose (2 g/kg, ip) or insulin (1.7 IU/kg, ip). Blood glucose was monitored via tail vein sampling using MediSure^®^ glucometers. The area under the curve (AUC) derived from GTT was used as an index of whole glucose excursion after glucose loading. Plasma glucose disappearance rate (Kitt) was calculated from the linear slope of the glucose concentration curve during ITT between 0 and 40 minutes.

### Glucose-stimulated insulin and incretin secretion (GSIS) tests

In figure 2, lean mice were transferred to clean cages without food and given a 5 mg/kg ip injection of the dipeptidyl peptidase 4 (DPP4) inhibitor Ile-Pro-Ile (Sigma, cat #I9759) followed by acute LPS challenge. Six hours later, blood was drawn from tail vein 30 minutes before and 15 minutes after oral glucose challenge (4g/kg) into tubes containing 10% (v/v) protease/DPP4 inhibitor cocktail (2-15 TIU of aprotinin/mL, 1.2 mg of EDTA/mL and 0.035 mg of Ile-Pro-Ile/mL). Figure 7a-i: following acute LPS stimulation in obese mice, animals were 6h fasted. Blood was collected from tail vein before (0 min) and after (20, 60 min) oral glucose challenge (4 g/kg). Blood glucose was monitored using MediSure^®^ glucometers, and blood samples were vortexed and then centrifuged (2000 g, 10 min, 4 °C) immediately. Plasma samples were stored at -80 °C for insulin and GLP-1 (7-36) amide determination using a multi-spot sandwich immunoassay kit (Meso Scale Discovery, cat #K15172C) or high sensitivity mouse insulin ELISA kit (Toronto Bioscience, Cat# 32270).

### Indices of insulin secretion and insulin resistance

Fasting insulin resistance was estimated using the homeostatic model assessment of insulin resistance (HOMA-IR) using the following formula (fasting insulin [mUI/mL] x fasting glucose [mmol/L]/22.5). Insulin resistance during the oral glucose load was estimated with the insulin resistance index (AUC^insulin^ x AUC^glucose^/100). β-cell function was estimated using the insulinogenic index, which was calculated as follows: ΔInsulin/ΔGlucose (pmol/mmol).

### *In vivo* gut permeability assay

Lean mice were chronically treated with different LPS and thereafter received 150 µL/mouse of a solution containing 80 mg of 4000 Da Fluorescein Isothiocyanate (FITC) Dextran/mL of saline. Blood samples were collected from tail vein using heparin coated capillary tubes before and 1 hour after gavage with FITC-Dextran. Plasma was obtained from blood samples by centrifugation (4°C, 8,000g, for 10 min), diluted in equal volume of PBS, and analyzed for FITC-dextran concentration with a fluorescence spectrophotometer (Excitation/Emission wavelengths 485/535 nm). Standard curves for calculating FITC-Dextran concentration were obtained by diluting FITC-dextran in PBS. Baseline readings were used to determine sample specific background fluorescence, which was subtracted from 1h readings. Circulating D-lactate was determined in baseline samples using PicoProbe D-Lactate Fluorimetric Assay Kit (Biovision).

### Intestinal Glucose Absorption

*In vivo* intestinal glucose absorption was measured 6 h following LPS challenge, where mice had access to water and no access to food. A non-metabolizable glucose analog (3-*O*-methyl-D-glucopyranose, 3-OMG, 4 mg/mouse) was gavaged to mice combined with a d-glucose (4 g/kg). 3-OMG quantified in the plasma by high-pressure liquid chromatography equipped with a triple quadrupole mass spectrometer. Briefly, deproteinized plasma samples were derivatized by acetylation and injected into an Agilent 1290 Infinity II HPLC with an Agilent 6495C iFunnel QQQ mass spectrometer. Analytes were separated on an Agilent RRHD Eclipse Plus C18 (100 mm x 2.1 mm, i.d., 1.8 μm) column using mobile phases consisting of (A) 0.1% (v/v) formic acid in water and (B) 0.1% (v/v) formic acid in acetonitrile and a constant flow of 0.4 mL/min. The autosampler and column were maintained at 10°C and 30 °C, respectively, throughout the analysis. The appearance of 3-OMG was monitored in the plasma over 90 min and the rate of intestinal glucose absorption was calculated using a linear regression model from time zero (i.e. before the gavage) to peak levels of 3-OMG.

### Messenger RNA extraction and RT qPCR

Total RNA was obtained from ∼30 mg of gonadal adipose tissue via mechanical homogenization at 4.5 m/s for 30 seconds using a FastPrep-24 tissue homogenizer (MP Biomedicals) and glass beads, followed by phenol–chloroform extraction. RNA was treated with DNase I (Thermo Fisher Scientific) and cDNA was prepared using 500 to 1000 ng of total RNA and SuperScript III Reverse Transcriptase (Thermo Fisher Scientific). Transcript expression was measured using TaqMan Assays with AmpliTaq Gold DNA polymerase (Thermo Fisher Scientific) and the ΔΔ*C*_T_ for target genes was calculated using *18S* housekeeping as a reference gene.

### Immunoblotting analysis

Analysis of total AKT and phospho (p)-AKT (serine 473) was performed as previously described (Steinberg et al., 2010). Mice were ip injected with saline or insulin (2.0 IU/kg) 15 min before euthanasia by cervical dislocation. Approximately 15 mg of livers were processed to yield total protein lysates, which were quantified using a BCA protein assay kit (Thermo Fisher Scientific). Twenty mg of protein were mixed with Laemmli buffer and loaded into 10% polyacrylamide gels. Samples were resolved by electrophoresis using a mini-Protean system (Bio-Rad) and transferred to polyvinylidene fluoride (PVDF) membranes, which were immunoblotted with antibodies for AKT (Cell Signaling, 1:1000) or pAKT (Cell Signaling, 1:1000) followed by incubation with HRP-linked anti-rabbit secondary antibody (Cell Signaling, 1:10000). Image documentation was carried out after incubation with Clarity Western ECL Substrate (Bio-Rad) in a ChemiDoc Imager (Bio-Rad).

## QUANTIFICATION AND STATISTICAL ANALYSIS

Data distribution was tested using Shapiro-Wilk test. For normally distributed data sets, unpaired t-test was used to compare two groups, and one-way analysis of variance (ANOVA) followed by Tukey’s multiple comparison test were used to compare between three or more groups. For non-parametric data sets, Mann-Whitney U test was applied to compare two groups, and Kruskal-Wallis followed by Dunn’s multiple comparisons tests were used to compare three or more groups. When time was considered as an additional variable, two-way repeated measures ANOVA with Tukey’s post hoc test was used for statistical testing. All graphs and statistical analyses were completed using GraphPad Prism 7. Data are represented as mean ±SEM, and the dispersion of each mouse within groups is represented by specific symbols. Statistical significance was accepted at *p* < 0.05, and all tests were two-sided.

## RESOURCE TABLE

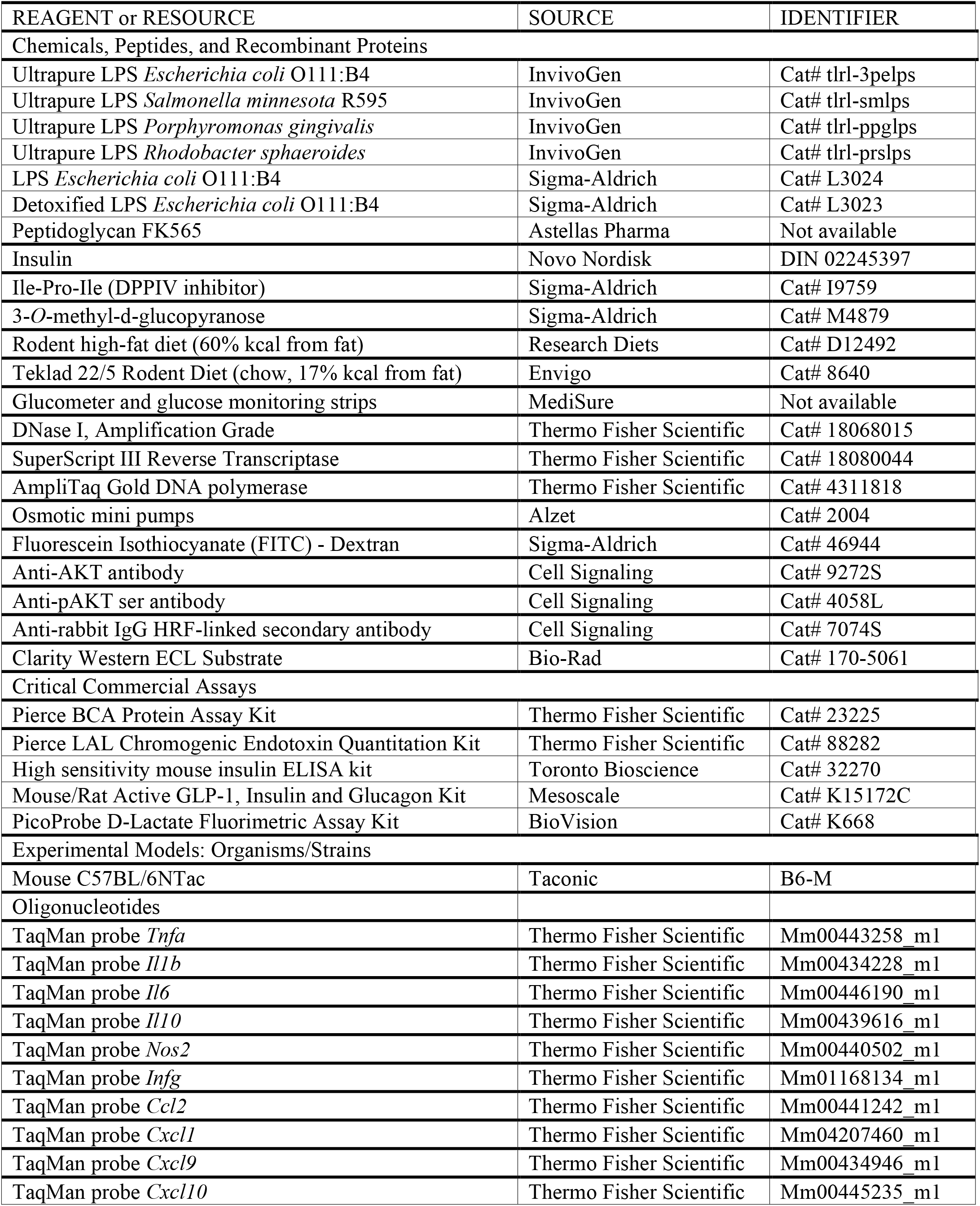
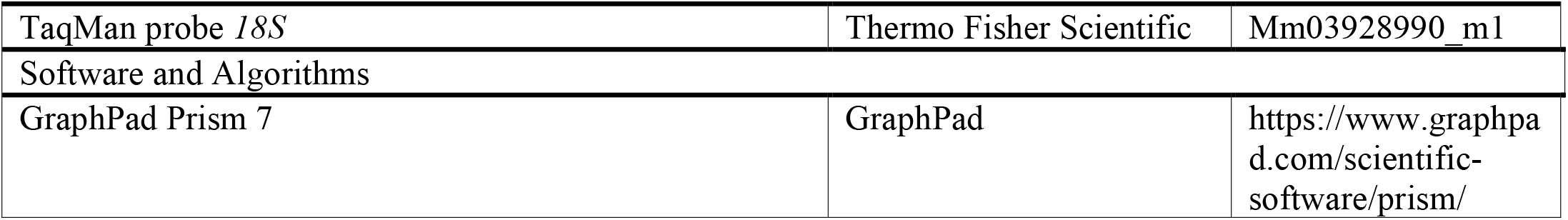

## Notes

### Competing Interest Statement

The authors have declared no competing interest.

